# Role of the hippocampus during logical reasoning and belief bias in aging

**DOI:** 10.1101/854208

**Authors:** Maryam Ziaei, Mohammad Reza Bonyadi, David C. Reutens

**Affiliations:** Centre for Advanced Imaging, the University of Queensland, Brisbane, Australia

**Keywords:** Hippocampus, aging, logical reasoning, white matter tract, cingulum, functional connectivity, multivariate analysis

## Abstract

Reasoning requires initial encoding of the semantic association between premises or assumptions, retrieval of these semantic associations from memory, and recombination of information to draw a logical conclusion. Currently-held beliefs can interfere with the content of the assumptions if not congruent and inhibited. This study aimed to investigate the role of the hippocampus and hippocampal networks during logical reasoning tasks in which the congruence between currently-held beliefs and assumptions varies. Participants of younger and older age completed a series of syllogistic reasoning tasks in which two premises and one conclusion were presented and they were required to decide if the conclusion logically followed the premises. The belief load of premises was manipulated to be either congruent or incongruent with currently-held beliefs. Our whole-brain results showed that older adults recruited the hippocampus during the premise integration stage more than their younger counterparts. Functional connectivity using a hippocampal seed revealed that older, but not younger, adults recruited a hippocampal network that included anterior cingulate and inferior frontal regions when premises were believable. Importantly, this network contributed to better performance in believable inferences, only in older adults group. Further analyses suggested that, in older adults group, the integrity of the left cingulum bundle was associated with the higher correct rejection of believable premises more than unbelievable ones. Using multimodal imaging, this study highlights the importance of the hippocampus during premise integration and supports the compensatory role of the hippocampal network during a logical reasoning task among older adults.

## Introduction

Logical reasoning, drawing a reasonable conclusion from related facts and assumptions, plays a central role in personal, complex political and societal decisions. Beliefs and prior knowledge, however, may contradict the given information and if they overshadow logic, unwarranted conclusions may be drawn; a phenomenon is known as belief bias (De Neys, 2012; Evans et al., 1983). In this situation, inhibition of currently-held beliefs is required to reach a logical conclusion, mainly engaging the prefrontal areas (Goel et al., 2000). While mounting evidence supports the role of frontal cortices in belief bias when currently-held beliefs contradict the given assumptions (for a review see Prado et al., 2010), the role of subcortical regions, such as the hippocampus, has not been fully understood. Indeed, there has been increasing evidence suggesting the importance of the hippocampus during various reasoning tasks. For instance, Goel et al. (2004) found that the hippocampus is involved during a reasoning task about the familiar spatial environment. Zeithamova and colleagues (2012) have also examined the role of the hippocampus in retrieving individual memories to answer a novel question in inferential reasoning. Another study reported that the hippocampus was active during a transitive inference, in which consideration of multiple relations is required to reach a logical conclusion (Wendelken & Bunge, 2010). While these studies have highlighted the importance of the hippocampus in reasoning tasks, the exact form of hippocampal engagement and its connection with the prefrontal areas in syllogistic reasoning has not been thoroughly investigated. Given the hippocampus involvement in the retrieval of semantic knowledge and in detecting conflicts between the current situation and prior experience (Kumaran & Maguire, 2007), it is reasonable to assume its role in syllogistic reasoning. This is particularly important during the assumption integration stage where given assumptions are compared with currently-held beliefs retrieved from the memory. We suggest that the hippocampus is essential for a flexible and valid reasoning decision and activity of the hippocampus, thus, it is expected to appropriately construct, manipulate, and update the information to respond to the task at hand.

### Age-related changes in the structure and function of the hippocampal-cortical networks

Existing literature indicates a complex pattern of activities in the aging brain during cognitive tasks. One view posits an over-recruitment of alternate brain circuits which is associated with maintained behavioral performance among older adults. In this view, increased activity of the prefrontal and other areas is interpreted as a compensatory mechanism (Davis et al., 2008). Contrary to the over-recruitment view, some studies found no difference between two age groups in performance or brain activity, supporting the brain maintenance hypothesis (for a review see Nyberg et al., 2012) and others found age-related decline (Nyberg et al., 2010; Persson et al., 2006). More specifically to underlying brain networks, a pattern of hippocampal connections with cortical areas, whether directly or indirectly, is altered in aging (For a detailed overview see Eichenbaum, 2017). Such a change in connectivity pattern of the hippocampus has been linked to changes in cognitive functions such as memory in late adulthood (Carr et al., 2017; Fjell et al., 2016; Salami et al., 2014; Samson & Barnes, 2013).

Despite reports of age-related hippocampus-prefrontal cortex (PFC) connectivity pattern alterations, little is known about changes in this connectivity during a complex cognitive task such as logical reasoning. To date, only a few studies have investigated age-related differences in belief bias and reasoning. De Neys and colleagues (2009) reported a decreased reasoning performance among older adults when belief and logic were in conflict, but not when they were congruent. In another study, Tsujii and colleagues (2010) replicated these findings and reported that older adults, unlike younger counterparts, recruited the bilateral inferior frontal gyrus (IFG) when belief and logic were in conflict. While these studies highlight the decline in reasoning performance among older adults, our understanding of neural networks underpinning a logical reasoning task in late adulthood is still incomplete. We believe that a complex, higher-order cognitive function such as reasoning undoubtedly relies on several brain structures and their connections. Thus, it is reasonable to suggest that these processes would be supported by a large-scale, interactive functional network, rather than by isolated brain regions such as IFG. Unlike the majority of previous studies that have focused on activity patterns of one single region, in the current study we have utilized a multivariate method to examine *functional connections* between areas during a reasoning task. Therefore, the primary aim of this study was to delineate age-related differences in functional engagement of the hippocampal network, and its strength of connectivity, as a function of believability load of syllogisms.

In addition to changes in the function of the hippocampus and its network as mentioned above, white matter integrity has shown to undergo substantial changes in aging and contribute to age-related cognitive decline (Madden et al., 2009). There is convincing evidence for age-related changes in some white matter microstructure that affect cognitive functions (Charlton et al., 2006; Choi et al., 2005; Fjell & Walhovd, 2010; Goh & Park, 2009; Nordahl et al., 2006; Tuch et al., 2005). Specifically, the integrity of white matter tracts, such as uncinate fasciculus and cingulum bundle, are integral for inhibitory control and executive functioning tasks (Catani, 2010; Grieve et al., 2007; Li et al., 2018). Changes in these tracts have been associated with an altered performance during executive function and inhibitory control in late adulthood (Davis et al., 2009; Hasan et al., 2009; Li et al., 2018; Vogt et al., 1992), demonstrating a possible link between their structural integrity and performance during a reasoning task. Our secondary aim was, therefore, to investigate whether the structural integrity of the white matter tracts, such as the cingulum bundle and uncinate fasciculus, contributes to changes in reasoning performance in aging.

### Current study

The aim of this study was twofold: first, to investigate age-related differences in hippocampal networks during a logical reasoning task, and second, to examine a relationship between the structural integrity of white matter tracts and logical reasoning performance across both age groups. Younger and older participants performed a syllogistic reasoning task in which they identified if a conclusion logically followed two given premises where the believability load of the premises was manipulated. Premises were either believable (congruent with currently-held belief: e.g. all parrots are birds), unbelievable (incongruent with currently-held beliefs: e.g. all lizards are mammals) or neutral (no believability load: all sothods are birds – where sothods is a pseudo-word). Three main analyses were conducted. First, a whole-brain analysis using task PLS was conducted to examine age-related differences in brain activity patterns as a function of the believability content of the syllogisms. Given the differences in analytical methods and lack of age group comparison in previous literature, there is a need to discern neural correlates underlying logical reasoning first to establish statistical support for further, connectivity-based, analyses. Second, a brain-behavior connectivity analysis using seed-behavioral PLS was performed to explore age-related differences in the hippocampal network as a function of believability load of assumptions. Importantly, we aimed to assess if the functional connectivity strength of the hippocampus was related to task performance during the syllogism task. Third, using structure-function analysis, we investigated whether the structural integrity of tracts involving the hippocampi, such as the cingulum bundle and the uncinate fasciculus, was correlated with logical reasoning performance.

In order to achieve these aims, we first identified brain regions that are critical for encoding the believability load of assumptions. A key region, such as the hippocampus, was chosen based on whole-brain findings and results of the previous meta-analysis on the importance of this region in memory and logical reasoning performance (Burgess et al., 2002; Goel et al., 2004; Zeithamova et al., 2012). First, we expected that the performance of younger and older adults during the logical reasoning task would be statistically different, youngers would perform better than the older participants. Subsequently, given previous evidence on the age-related differences in hippocampal functional connectivity, we hypothesized that younger and older adults would show differential recruitment of hippocampal brain networks in response to believable and unbelievable assumptions, contributing to differences in behavioral performance during a logical reasoning task. Alternatively, if younger and older adults show a lack of difference in hippocampal functional network engagement, results from functional connectivity should reveal a common network between two age groups. This prediction is evaluated by the seed-behavioral PLS which tests whether the networks connected to the hippocampus are different between two age groups as a function of believability load. Additionally, we examined age-related differences in the white matter structural integrity, measured by the fractional anisotropy (FA), between two age groups and tested whether such difference would contribute to differences in behavioral performance during a logical reasoning task. This was evaluated by structure-behavior analysis.

## Material & Methods

### Participants

Thirty-one healthy younger and thirty-two healthy older adults participated in this study. Due to extensive head movement and brain signal loss, two older and two younger adults were excluded from the analysis, leaving 29 younger adults (aged 18-26 years; *M* = 21.13, *SD* = 2.72; 15 females) and 30 older adults (aged 61-78 years; *M* = 70.34, *SD* = 4.27; 15 females). Younger adults were recruited from The University of Queensland and were reimbursed either with course credits or AUD$15 per hour. Older adults were volunteers from the community recruited through flyers on notice boards in local Rotary clubs, University of Third Age, libraries, churches, and The University of Queensland’s Aging Mind Initiative. Older adults were reimbursed AUD$20 per hour. Participants were screened for MRI compatibilities, mood disorder (depression and anxiety), claustrophobia, significant neurological and psychiatric disorders such as epilepsy and head injury before enrolment in the study. All participants were English speakers, right-handed, with normal or corrected-to-normal vision using MRI compatible glasses. Older adults underwent additional screening to rule out cognitive decline on the Mini Mental State Examination (Folstein et al., 1975), a widely used dementia screen; all older adults scored above the recommended cut-off of 24 (*M* = 29.34, *SD* = 0.82). All participants took part in two separate test sessions, the first involving MRI scanning and the second involving behavioral and neuropsychological assessments. All participants were provided with written consent forms and were debriefed upon the completion of the second session. The experiment was approved by the Bellberry Human Research Ethics Committee.

### Task design and materials

A logical *statement* in this study follows a generic form of <*quantifier, subject, copula, predicate*>, e.g., All dogs are animals, where “All” is a quantifier, “dogs” is a subject, “are” is a copula, and “animals” is a predicate. Logical arguments in this study were in the form of standard syllogisms and included three statements: two premises and one conclusion. The subject and the predicate of a premise was formed by arbitrary sets (e.g., dogs, mammals, furniture). The two premises had exactly one set in common that may appear in either the subject or the predicate in either of the premises (*Set*_*2*_ in Table 1). Hence, the two premises involved exactly three sets (*Set*_*1*_, *Set*_*2*_, and *Set*_*3*_ in the example in Table 1), two of which are uniquely used in each premise and one which is used in both. The conclusion of a syllogism provides a statement about the sets that appear uniquely in premises (*Set*_*1*_ and *Set*_*3*_ in the example). A conclusion “follows” from the premises if the premises provide conclusive evidence to support it. Otherwise, the conclusion “does not follow” from the premises, either because the conclusion is wrong given the premises, or is not completely supported by the premises. A full description of the task is presented in Ziaei et al. (2019).

**Table 1.** Form of syllogisms, quantifiers, subjects, copula, and predicate used in the experimental design

|  |  | Quan | S | C | Pre |
| --- | --- | --- | --- | --- | --- |
|  | tifier |  | ubject | opula | dicate |
| mise 1 | Pre | All | Set<sub>1</sub> | is | Set<sub>2</sub> |
| mise 2 | Pre | No | Set<sub>2</sub> | is | Set<sub>3</sub> |
| clusion | Con<br>e | Som | Set<sub>3</sub> | is<br>not | Set<sub>1</sub> |

Additionally, in a syllogism, each statement includes a quantifier (All, No, and Some) and a copula (Is or Is not). We used propositions A (All, is) and E (No, is) on all premises (counterbalanced across all runs). Given that proposition E simplifies the reasoning task substantially, we avoided the use of proposition E in both premises. For the conclusions, however, we used the appropriate propositions required to ensure that conclusions that followed or did not follow the premises were balanced.

The order of the sets and predicate of the premises was balanced. In this experiment, we used two types of ordering: i) the common set belonged to the subject of premise 1 and the subject of premise 2, and ii) the common set belonged to the predicate of premise 1 and the subject of premise 2. The conclusion takes the unique sets from each premise and combines them through a proposition. The logical reasoning task was to decide if the generated statement follows from the premises or not. Whether the subject of the conclusion was drawn from the first or second premise was also counterbalanced within runs.

Conclusions and premises were either believable or unbelievable in terms of the belief load. For example, “all dogs are animals” is a believable statement while “all birds are mammals” is an unbelievable statement. In addition, control premises with a neutral load comprising a meaningless pseudo-word were used (e.g. “all parrots are nickhomes”, where “nickhomes” is a neutral word without any belief load). Pseudo-words were only used on the premises so that they were a shared set between the premises. Hence, the first premise was always believable, the second premise’s believability was manipulated (believable, unbelievable, or neutral), while the conclusion was either believable or unbelievable. The followings are two examples of syllogisms with a believable premise / unbelievable conclusion and an unbelievable premise / believable conclusion, respectively:

All pines are trees; No pines are willows; Therefore, all trees are willows (believable premise/unbelievable conclusion; logically invalid)

All lories are parrots; No parrots are animals; Therefore, some animals are not lories (unbelievable premise/believable conclusion; logically valid)

A total of 96 syllogisms were generated using an in-house algorithm. Six conditions included in the task: 1. believable premise/believable conclusion, 2. believable premise/unbelievable conclusion, 3. unbelievable premise/believable conclusion, 4. unbelievable premise/unbelievable conclusion, 5. neutral premise/believable conclusion, 6. neutral premise/unbelievable conclusion (see Supplementary Material for access to all of the syllogisms used in the study).

### Experimental design

Before the scan, participants were instructed about the task and procedure of the scanning session. A practice run was administered until they were familiar with the timing and instruction of the task. The imaging session included two components: two structural MRI scans (T1-weighted scans and Diffusion-Weighted Imaging (DWI) scans) and the logical reasoning task with functional MRI (fMRI), all lasted for 45 minutes in total. During the logical reasoning task, participants were asked to determine if the conclusion statement logically followed from the two premises using two keys on an MRI-compatible response box. The first premise was presented for 2 seconds followed by a second premise for 4 seconds. After the second premise, the conclusion statement was presented for 12 seconds (Figure 1). All statements (premises and conclusion) remained on the screen until the end of the presentation of the conclusion to reduce the working memory load. A jittered fixation cross was presented after the conclusion with four-time intervals: 0.5 seconds (24 trials), 1 second (24 trials), 1.5 seconds (24 trials), and 2 seconds (24 trials). The task consisted of 6 runs, each run lasting for 5.16 minutes with three runs of the task presented before and three after the structural scan.

**Figure 1.**
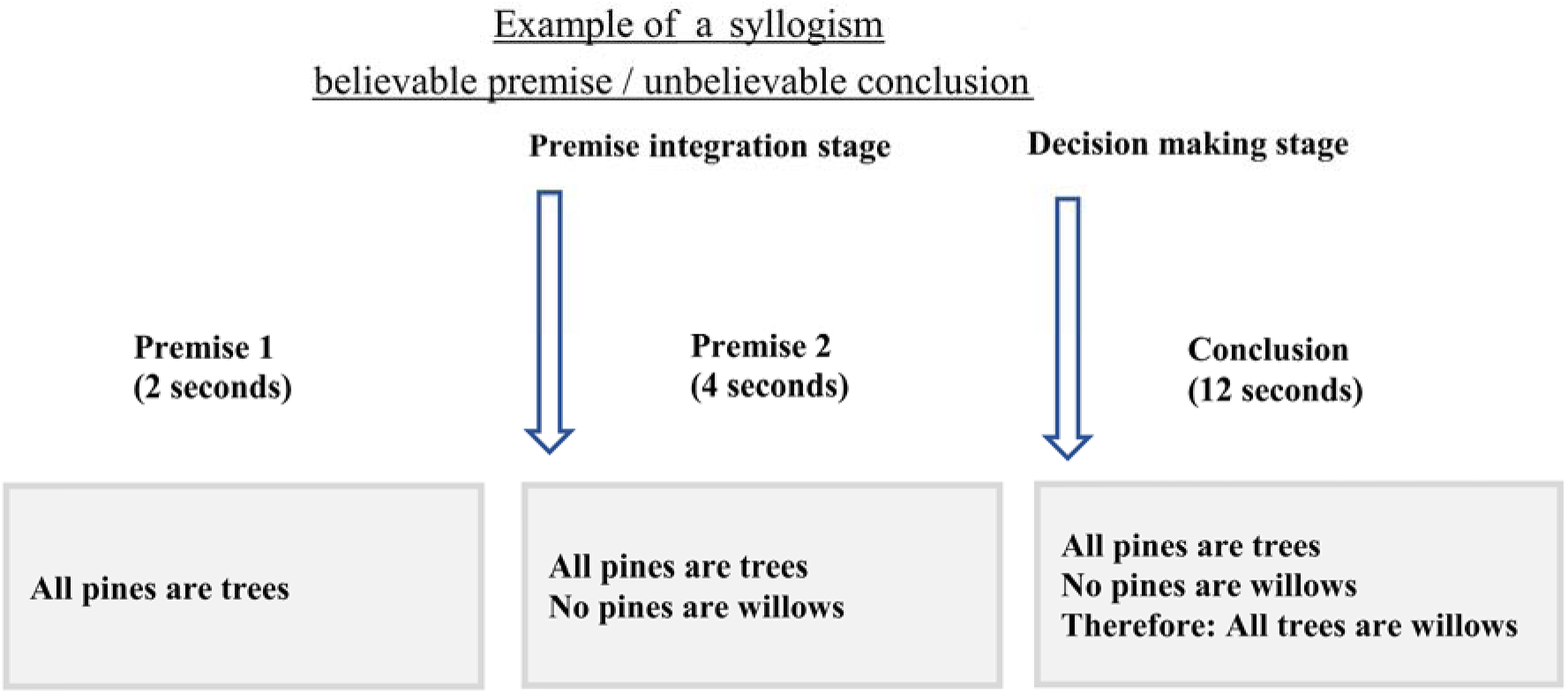
The timing and example of the experimental design. In this experiment, participants were presented with the first premise for two seconds followed by the second premise that was shown for 4 seconds. During the conclusion presentation, 12 seconds, participants were asked to choose if the conclusion follows the two premises or not, using the MRI compatible response box. Onsets from the second premise were only used to assess brain activity during premise integration.

### Background measures

In addition to the imaging session, all participants completed several tasks to assess emotional well-being as measured by the Depression, Anxiety, Stress Scale (DASS-21; Lovibond & Lovibond, 1995), executive functioning as measured by the Stroop task (Jensen & Rohwer, 1966) and the Trail Making Test (Reitan & Wolfson, 1986), and intelligence as measured by the National Adult Reading Test (Nelson, 1982). Descriptive and inferential statistics of background measures are reported in Table 2.

**Table 2.**
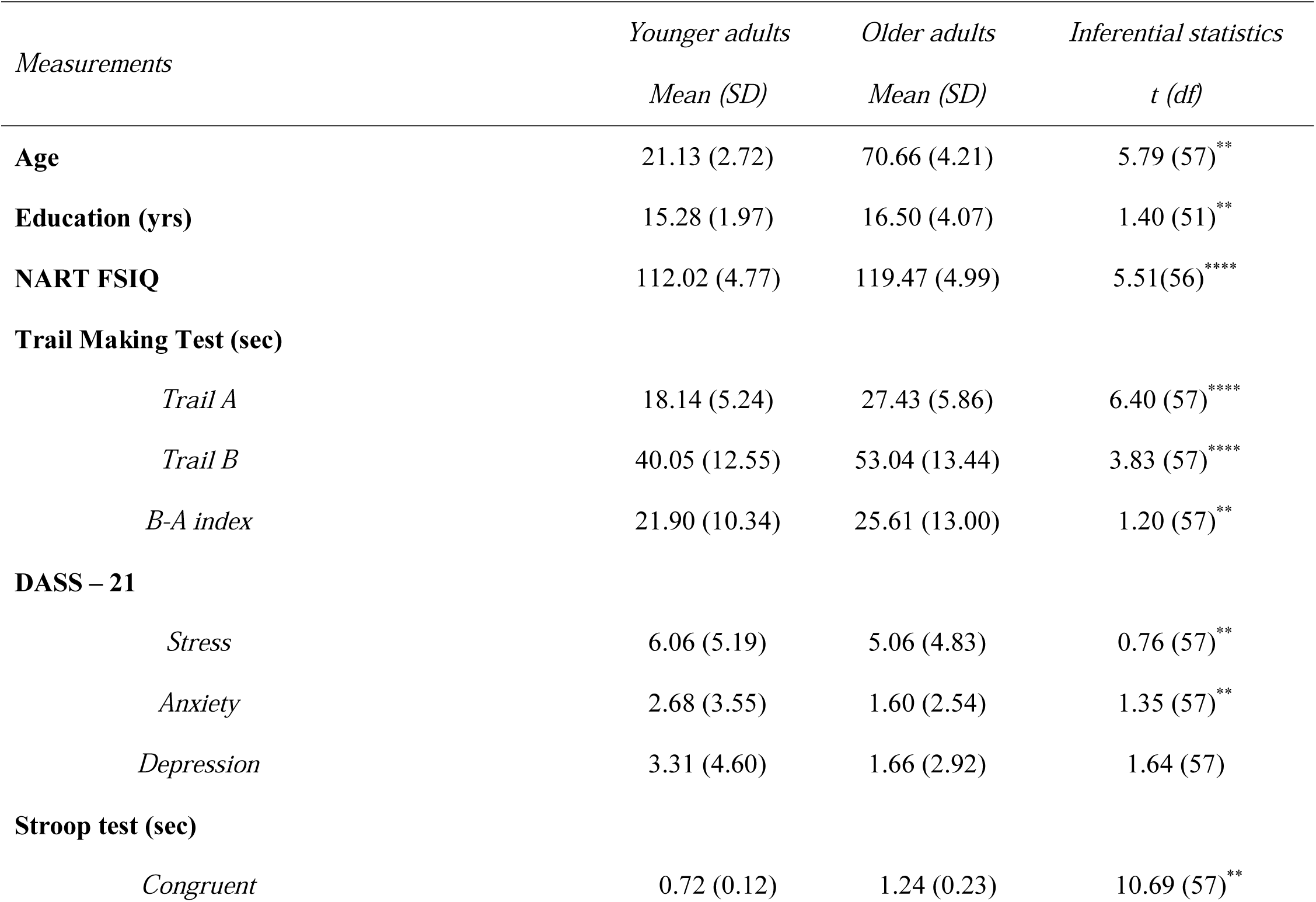
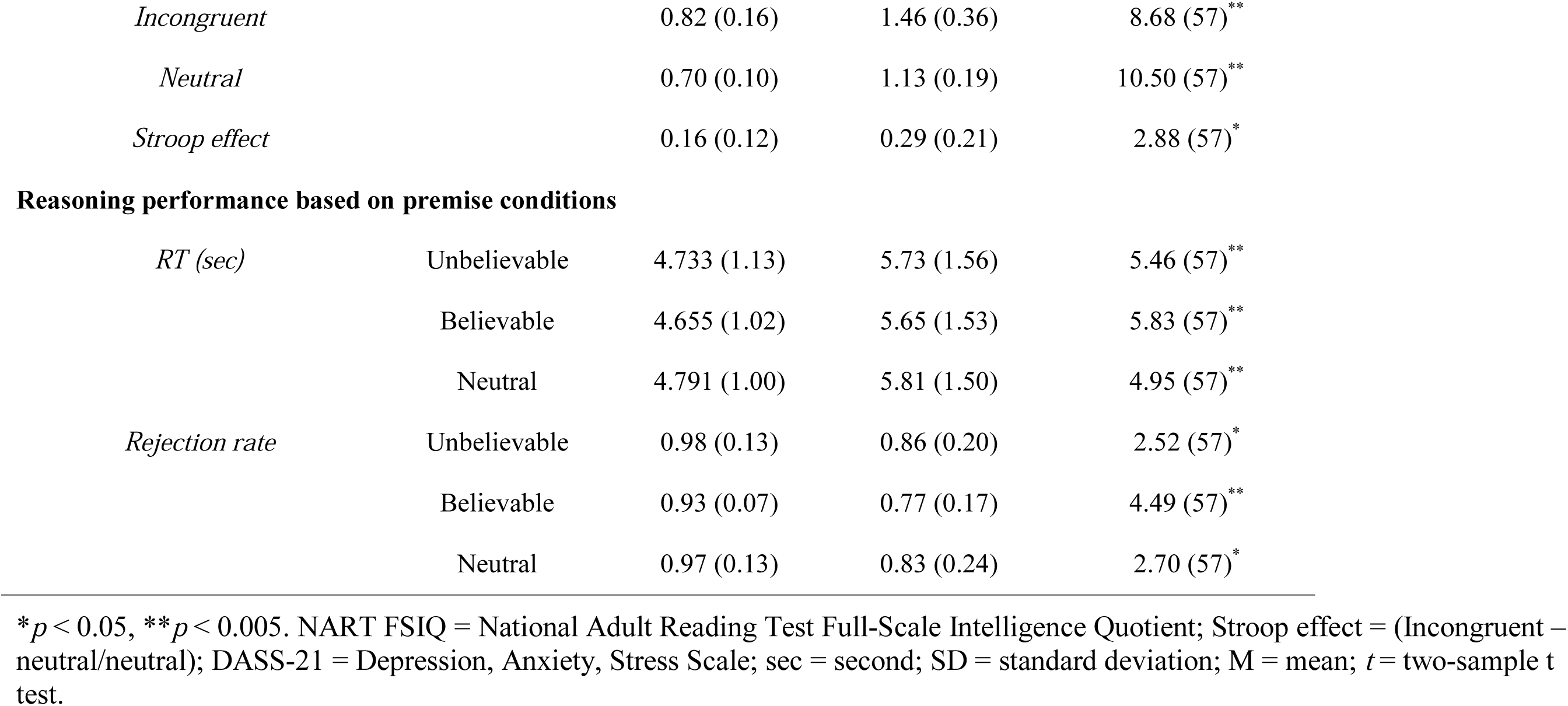
Descriptive and inferential statistics of performance on background measures and logical reasoning task

| *Measurements* | *Younger adults* | *Older adults* | *Inferential statistics* |
| --- | --- | --- | --- |
|  | *Mean (SD)* | *Mean (SD)* | *t (df)* |
| **Age** | 21.13 (2.72) | 70.66 (4.21) | 5.79 (57)** |
| **Education (yrs)** | 15.28 (1.97) | 16.50 (4.07) | 1.40 (51)** |
| **NART FSIQ** | 112.02 (4.77) | 119.47 (4.99) | 5.51(56)**** |
| **Trail Making Test (sec)** |  |  |  |
| *Trail A* | 18.14 (5.24) | 27.43 (5.86) | 6.40 (57)**** |
| *Trail B* | 40.05 (12.55) | 53.04 (13.44) | 3.83 (57)**** |
| *B-A index* | 21.90 (10.34) | 25.61 (13.00) | 1.20 (57)** |
| **DASS – 21** |  |  |  |
| *Stress* | 6.06 (5.19) | 5.06 (4.83) | 0.76 (57)** |
| *Anxiety* | 2.68 (3.55) | 1.60 (2.54) | 1.35 (57)** |
| *Depression* | 3.31 (4.60) | 1.66 (2.92) | 1.64 (57) |
| **Stroop test (sec)** |  |  |  |
| *Congruent* | 0.72 (0.12) | 1.24 (0.23) | 10.69 (57)** |
| *Incongruent* | 0.82 (0.16) | 1.46 (0.36) | 8.68 (57)** |
| *Neutral* | 0.70 (0.10) | 1.13 (0.19) | 10.50 (57)** |
| *Stroop effect* | 0.16 (0.12) | 0.29 (0.21) | 2.88 (57)* |

**Reasoning performance based on premise conditions**
|  |  |  |  |  |
| --- | --- | --- | --- | --- |
| *RT (sec)* | Unbelievable | 4.733 (1.13) | 5.73 (1.56) | 5.46 (57)** |
|  | Believable | 4.655 (1.02) | 5.65 (1.53) | 5.83 (57)** |
|  | Neutral | 4.791 (1.00) | 5.81 (1.50) | 4.95 (57)** |
| *Rejection rate* | Unbelievable | 0.98 (0.13) | 0.86 (0.20) | 2.52 (57)* |
|  | Believable | 0.93 (0.07) | 0.77 (0.17) | 4.49 (57)** |
|  | Neutral | 0.97 (0.13) | 0.83 (0.24) | 2.70 (57)* |

\* $p < 0.05$ , \*\* $p < 0.005$ . NART FSIQ = National Adult Reading Test Full-Scale Intelligence Quotient; Stroop effect = (Incongruent – neutral/neutral); DASS-21 = Depression, Anxiety, Stress Scale; sec = second; SD = standard deviation; M = mean; $t$ = two-sample $t$ test.

### Image acquisition

Functional images were acquired at the Centre for Advanced Imaging using a 3T Siemens scanner with a 32-channel head coil. The functional images were obtained using a whole-head T2*-weighted multiband sequence (473 interleaved slices, repetition time (TR) = 655ms, echo time (TE) = 30ms, voxel size = 2.5mm^3^, field of view (FOV) = 190mm, flip angle = 60º, multi-band acceleration factor = 4). High-resolution T1-weighted images were acquired with an MP2RAGE sequence (176 slices with 1mm thickness, TR = 4000ms, TE = 2.89ms, voxel size = 1mm^3^, TI = 700ms, FOV = 256mm). Participants were provided with cushions and earplugs around their head inside the head coil to minimize the noise and head movement. Participants observed the task on a computer screen through a mirror mounted on top of the head coil.

### fMRI Preprocessing

For functional analysis, T2*-weighted images were preprocessed with Statistical Parametric Mapping Software (SPM12; http://www.fil.ion.ucl.ac.uk/spm) implemented in MATLAB 2015b (Mathworks Inc., MA). Following the realignment to a mean image for head-motion correction, images were segmented into gray matter, white matter, and cerebrospinal fluid. Then, images were spatially normalized into a standard stereotaxic space with a voxel size of 2 mm^3^, using the Montreal Neurological Institute (MNI) template, and then spatially smoothed with a 6 mm^3^ Gaussian Kernel. None of the participants included in the analyses had head movement above 1mm.

### fMRI analyses

The imaging data were analyzed using a multivariate analytical method Partial Least Squares analysis (PLS; McIntosh et al. (1996)) as implemented in PLS software running on MATLAB (The MathWorks Inc., MA). For a detailed tutorial and review of PLS, see Krishnan et al. (2011). PLS analysis uses singular value decomposition (SVD) of a single matrix that contains all data from all participants to find a set of orthogonal latent variables, which represent a linear combination of the original variables. PLS decomposes all images into a set of patterns that capture the greatest amount of covariance in the data, without making assumptions about conditions or imposing contrasts. Using PLS enables us to differentiate contribution of different brain regions in relation to the task demands, activation of a functional seed, behavioral or anatomical covariates. Each latent variable delineates cohesive patterns of brain activity related to experimental conditions. Usually, the first latent variable accounts for the largest covariance of the data and progressively smaller amounts of covariance are attributed to subsequent latent variables. The brain score reflects how much each participant contributes to the pattern expressed in each latent variable. Therefore, each latent variable consists of a singular image of voxel saliences (*i.e*., a spatiotemporal pattern of brain activity), a singular profile of task saliences (*i.e*., a set of weights that indicate how brain activity in the singular image is related to the experimental conditions, functional seeds, or behavioral/anatomical covariates), and a singular value (*i.e*., the amount of covariance accounted for by the latent variable). There is no need for multiple comparison correction, as the activation patterns identified by PLS and corresponding brain responses are done in a single mathematical step (McIntosh et al., 2004).

The statistical significance of each latent variable was assessed using a permutation test, which determines the probability of a singular value from 500 random reorderings (McIntosh et al., 1996). Additionally, to determine the reliability of the saliences for each brain voxel, the standard error of each voxel’s salience on each latent variable was estimated by 100 bootstrap resampling steps (Efron & Tibshirani, 1985). Peak voxels with a bootstrap ratio (*i.e.*, salience/standard error) > 3 were considered to be reliable, as this approximates *p* < 0.001 (Sampson et al., 1989).

In the current study, we used two independent analyses: task PLS and seed-behavioral PLS. Given this is the first study of its kind that examines age-related differences in syllogisms where the believability load of premises is manipulated, first, we aimed to determine the whole-brain activity pattern during the assumption integration stage (second premise) as a function of believability load of statements across both age groups. Second, we assessed age-related differences in the hippocampal functional network to examine whether the strength of activity in the hippocampal network is modulated by the believability load of the second premise and whether the hippocampal network contributes to the behavioral performance. We aimed to explore the neural correlates of age-related differences during the premise integration stage, our fMRI analyses focused on the second premise’s believability load, collapsing across the believability load of the conclusion.

### Whole-brain analysis (Task PLS)

The whole-brain analysis focused on the onset times from the beginning of the *second* premise and included all three belief loads, depicted in Figure 2. Given that the logical reasoning task in this study was event-related, the activity at each time point in the analysis was normalized to activity in the first TR from the second premise. Using this approach, we examined if the neural correlates for believable, unbelievable or neutral premises were different between younger and older adults. Thus, both age groups and three premise conditions were included in the analyses, simultaneously. This analysis revealed two latent variables and for clarity, we depicted the results of each LV and their positive/negative saliences in separate panels.

**Figure 2.**
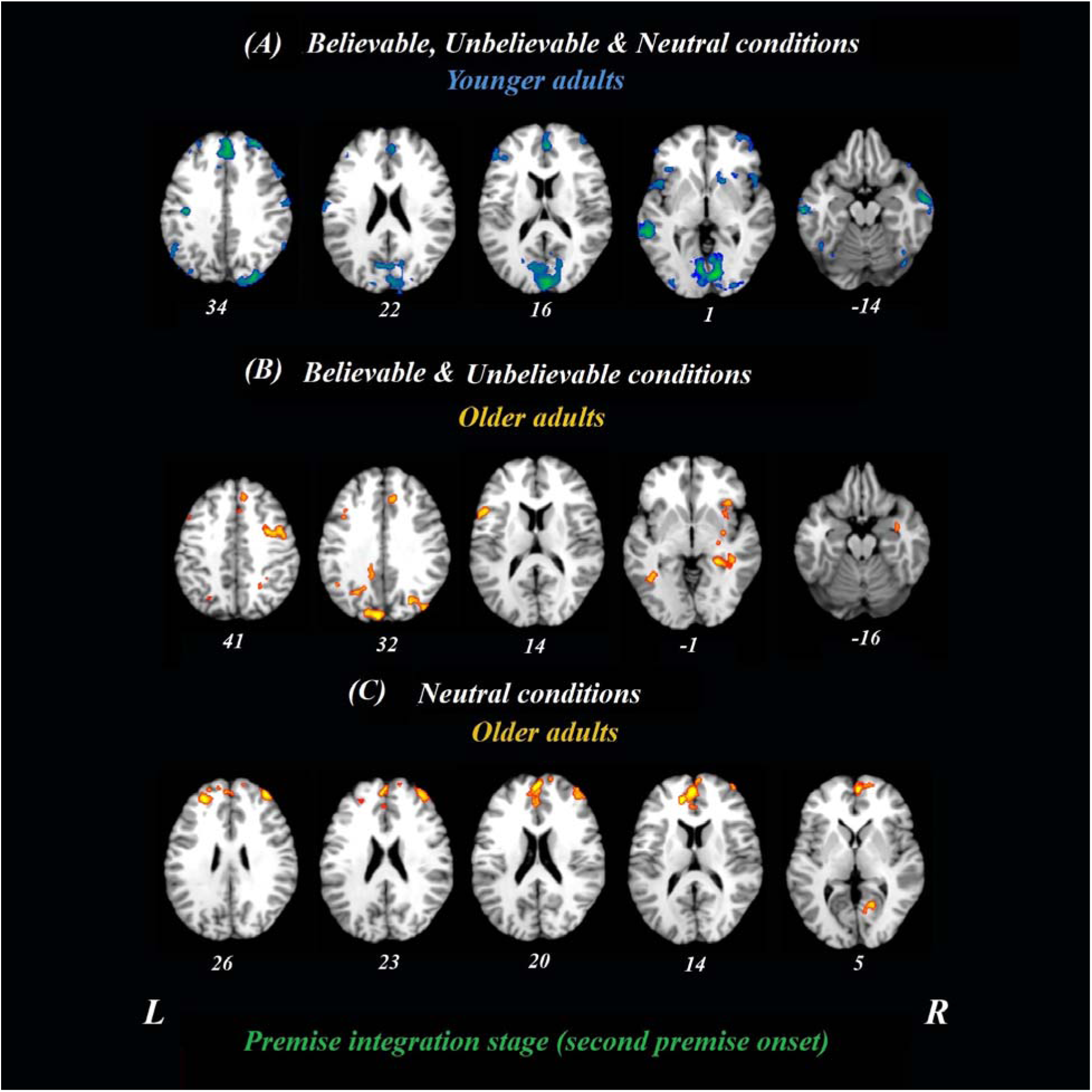
Whole-brain results during premise integration stage in both age groups. Onsets of the second premise were used in this analysis using task PLS. All believable, unbelievable, and neutral premises were included in this analysis. ***(A)***, Younger adults recruited the blue regions similarly for all conditions during the premise stage. ***(B)***, Older adults, however, recruited these yellow regions more during unbelievable and believable premises. ***(C)***, these regions were more involved during neutral premises relative to other conditions among older adults. All reported regions have bootstrap ratio of ≥ 2.5 and cluster size ≥ 50 voxels. Abbreviations: L = left hemisphere, R = right hemisphere.

### Brain-behavior connectivity analyses (Seed-behavioral PLS)

We also examined task-related functional connectivity for the hippocampal seed region and assessed the relationship with behavioral performance. To delineate the functional network involved during the premise integration stage, signal intensity values from peak voxels of the seed were extracted and correlated with activity in the rest of the brain, as well as behavioral performance across all participants for believable and unbelievable premises. Our peak voxel (28-37 −2) was selected based on its activity from our whole-brain analysis and previous functional studies reported in the Neurosynth dataset (Z score of 11.22; Yarkoni et al., 2011). In an independent analysis of the task PLS, these correlations were then combined into a matrix and decomposed with singular value decomposition in a separate and independent analysis than whole-brain analysis. The seed-behavioral PLS analysis calculates a set of latent variables characterizing a set of regions for which the activity was correlated with the seed region, the hippocampus, and with the behavioral performance during both believable and unbelievable premises. Permutation and bootstrap sampling were used to determine the significance and reliability of the functional connectivity analyses as in the whole-brain analysis.

### DWI acquisition and analysis

We used the FMRIB’s Diffusion Toolbox (FDT) (Andersson & Sotiropoulos, 2016) to correct our DWI images for eddy current distortion and head motion. Fractional Anisotropy (FA) in each voxel was estimated by first removing the non-brain tissues from the corrected images using the Brain Extraction Tool (Smith, 2002) and then locally fitting the diffusion tensor model at each voxel using the FDT. FA is a marker of the integrity of white matter tracts, reflecting the coherence within a voxel and fiber density (Alexander et al., 2007; Beaulieu, 2002). FMRIB’s Linear Image Registration Tool (FLIRT) (Greve & Fischl, 2009; Jenkinson et al., 2002; Jenkinson & Smith, 2001), with 12 degrees of freedom and trilinear interpolation, was used to realign the FA map of each subject with the standard brain template, MNI152 T1 1mm isotropic voxels. The affine transformation provided by this procedure ensures the transformed FA maps are in the same 3D coordinate system. We then generated the desired white matter mask for the cingulum bundle and the uncinate fasciculus using the ICBM-DTI-81 white-matter atlas (Hua et al., 2008; Oishi et al., 2010; Wakana et al., 2007). The average FA value in each tract was calculated and used in the structure-behavior analysis.

### Structure-behavior analysis

Spearman correlation analyses with white matter structural integrity, FA values, and behavioral performance, the rejection rate, was conducted for each experimental condition and each age group separately. FA values of the cingulum bundle and uncinate fasciculus were included in the analyses. All behavioral analyses were performed in IBM SPSS statistics (version 25) with the *p-*value of 0.05 to be the significance level.

#### Statistical analysis of behavioral data

Hence, a high performance refers to a high rejection rate in believable statements and a low rejection rate in unbelievable statements. The rejection rate for each participant was defined as the number of rejected syllogisms divided by the total number of syllogisms that should have been rejected. Repeated measures ANOVA was performed for reaction times (RTs) and rejection rate as dependent variables. RTs were defined as the response times that took participants to decide during the conclusion stage. A 3 (premise belief load; believable, unbelievable, and neutral) by 2 (age group; younger and older adults) repeated measures ANOVA was conducted on RTs and rejection rate. Additional analyses on the reaction times and rejection rate of the conclusion stage have been reported in the Supplementary Results. Behavioral results considered statistically significant at a *p-*value of 0.05. No difference was found between males and females in the behavioral performance. Therefore, all analyses were conducted across both genders.

## Results

### Behavioral results

#### Rejection rate

Repeated measure ANOVA on rejection rate revealed a significant main effect of premise belief load (*F*(2,114) = 5.01, *p* = 0.008, 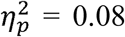), suggesting that believable premises had lower rejection rate relative to unbelievable and neutral ones (*t*(58) = 2.95, *p* = 0.004, *d* = 0.77 and *t*(58) = 2.32, *p* = 0.02, *d* = 0.60, respectively). A significant main effect of age group (*F*(1,57) = 14.19, *p* < 0.001, 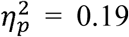) was found, suggested that older adults had lower rejection rate relative to younger adults in all conditions. However, the interaction between age group and premise belief load did not reach significance (*F*(2,114) = 0.40, *p* = 0.67, 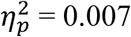).

#### Reaction Times

Repeated measures ANOVA on reaction times showed non-significant main effect of premise belief load (*F*(2,114) = 2.16, *p* = 0.12, 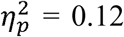), age group (*F*(1,57) = 31.83, *p* = 0.000, 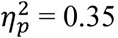), and interaction between age group and premise belief load (*F*(2,114) = 1.12, *p* = 0.32, 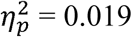).

#### FA values

There were significant differences between younger and older adults in both left (*t*(56) = 5.46, *p* < 0.001) and right (*t*(56) = 3.42, *p* = 0.001) uncinate fasciculus FA values, suggesting that older adults had lower FA values in this tract (M _left_ = 0.25, SD = 0.03; M _right_ = 0.26, SD = 0.04). relative to younger adults (M _left_ = 0.30, SD = 0.02; M _right_ = 0.29, SD = 0.02). There were no age-related differences in the FA values of the cingulum bundle (left: *t*(56) = 0.95, *p* = 0.34; Right: *t*(56) = 0.19, *p* = 0.84).

## fMRI results

### Whole-brain analysis (Task PLS)

#### Age-related differences between believable and unbelievable premises

To determine age-related differences in believable and unbelievable premises, the whole-brain results were conducted on the premise integration stage and revealed two latent variables. The first accounted for 68% of the covariance of the data (*p* < 0.001) and included right superior frontal gyrus, right anterior cingulate cortex (ACC), bilateral inferior frontal gyrus (IFG), left insula, left inferior parietal lobe, left precuneus, and left caudate. These regions were recruited by younger adults only, irrespective of the belief load of the premise (Table 3).

**Table 3.** Whole-brain results showing regions activated for all conditions among younger adults during the premise integration stage

| Regions | Hem | Cluster size | MNI coordinates | BSR |
| --- | --- | --- | --- | --- |
|  |  |  | XYZ |  |
| *All conditions – premise integration stage* |  |  |  |  |
| Superior frontal gyrus | R | 19027 | 16 34 54 | -8.0285 |
| Medial cingulate cortex | L | 117 | -4 -20 36 | -4.2812 |
| Middle cingulate cortex | R | 104 | 14 -42 36 | -4.3166 |
| Inferior frontal gyrus (p. Triangularis) | L | 381 | -48 40 14 | -5.1408 |
| Inferior frontal gyrus (p. Opercularis) | R | 232 | 50 12 0 | -4.9558 |
| Middle orbital gyrus | R | 60 | 6 32 -10 | -5.1013 |
| Superior medial gyrus | L | 16156 | -2 32 52 | -8.5392 |
|  | R | 180 | 0 54 6 | -5.1678 |
| Insula | L | 603 | -28 22 8 | -6.2758 |
| Superior temporal gyrus | L | 70 | -62 -46 16 | -4.0708 |
| Middle temporal gyrus | L | 785 | -58 -38 2 | -6.0463 |
|  | R | 564 | 58 -10 -18 | -5.5045 |
| Medial temporal gyrus | L | 857 | -56 -40 2 | -6.953 |
|  | R | 787 | 66 -14 -12 | -5.2644 |
| Inferior parietal lobule | L | 250 | -42 -80 26 | -5.336 |
| Thalamus | R | 59 | 16 -14 -4 | -5.4105 |
| Putamen | R | 544 | 16 12 -2 | -5.7981 |
| Caudate | L | 51 | -20 0 22 | -4.8374 |
| Precuneus | L | 81 | -14 -52 54 | -4.29 |
| Cerebellum | R | 92 | 22 -52 -18 | -4.9426 |

Hem = hemisphere; BSR = bootstrap ratio; MNI = Montreal Neurological Institute.

The second latent variable accounted for 10% of the covariance of the data and distinguished neutral conditions from believable/unbelievable conditions in older adults only (*p* = 0.048). One network included the left IFG, right insula, right middle frontal gyrus, left posterior cingulate cortex, and right hippocampus. These regions showed enhanced activity among older adults group during both believable and unbelievable premises more than neutral ones (Figure 2). Older adults also recruited a distinct sets of regions including right ACC, right superior frontal gyrus, and left middle frontal gyrus for neutral premises relative to other conditions (Table 4).

**Table 4.**
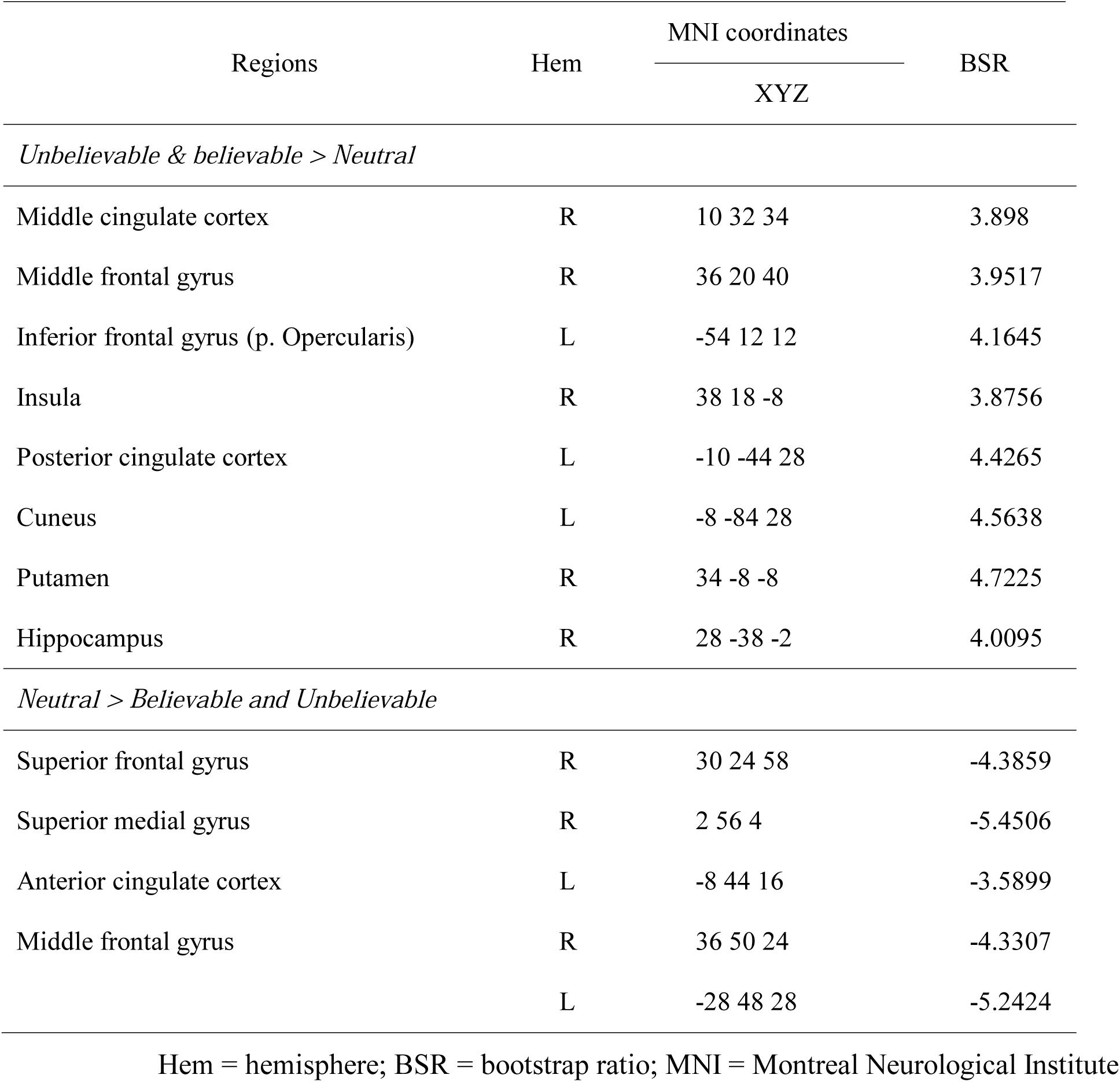
Whole-brain results showing regions modulated by the belief among older adults during the premise integration stage

### Brain-behavior connectivity results (Seed-behavioral PLS)

#### Age-related differences in the hippocampal functional network during premise integration

The brain-behavior connectivity analyses using the hippocampus as a seed with behavioral performance were conducted and revealed two significant latent variables. The first latent variable accounted for 26% of the covariance of the data (*p* < 0.001) and yield a network that was connected to the hippocampus for both believable and unbelievable conditions among younger adults without contributing to the performance. This hippocampal functional network contributed to better behavioral performance – rejection rate – among older adults during the believable conditions. This network included anterior cingulate, bilateral postcentral gyrus, right middle frontal gyrus, right superior parietal lobe, and right precuneus.

The second latent variable accounted for 14% of the covariance of the data (*p* = 0.042) and yield a network, which included anterior cingulate, left inferior frontal gyrus, bilateral insula, left superior temporal gyrus, bilateral precentral gyrus, left angular gyrus, left parahippocampus, left hippocampus, bilateral thalamus, posterior cingulate gyrus, precuneus, and bilateral lingual gyrus regions (Figure 3). This network was engaged by older adults only during the believable conditions and contributed to a higher rejection rate for believable conditions among this age group.

**Figure 3.**
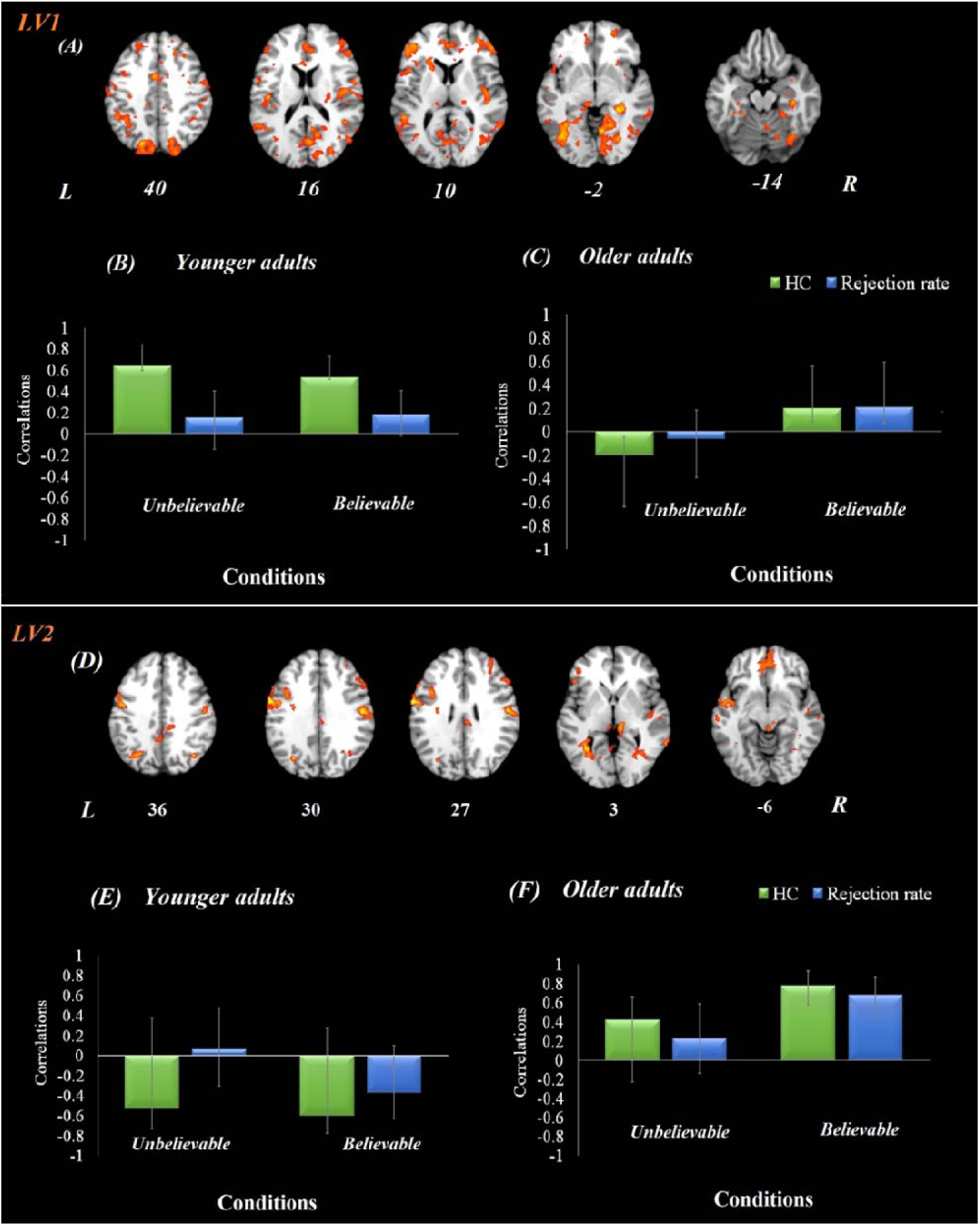
Hippocampal functional connectivity results in both age groups. Onsets of the premise stage were used for this analysis using seed-behavioral PLS. LV1 represents functional connectivity of hippocampal activity in the first latent variable. ***(A)*** The functional network connected to the hippocampus during the premise integration stage. Panel ***(B)*** and ***(C)*** show correlations between activity in the hippocampal brain network and behavioral performance among younger and older adults, respectively, for each experimental condition. LV2 represents functional connectivity of hippocampal activity in second latent variable, demonstrating compensatory network. Panel ***(D)*** demonstrates the functional network connected to the hippocampus during the premise integration stage. Panel ***(E)*** and ***(F)*** show correlations between activity in the hippocampal brain network and behavioral performance among younger and older adults, respectively, for each experimental condition For visualization purposes, bootstrap ratio threshold is set at 2.5 (*p* < 0.005); however, reported whole-brain activity is set at BSR 3 (*p* < 0.001). Error bars represent 95% confidence intervals, when crossing zero interprets as unreliable. Abbreviations: LV: latent variable, HC = hippocampus, L = left hemisphere, R = right hemisphere.

### Structure-behavior results

#### Age-related differences in the structural integrity of cingulum bundle for premise integration

To provide additional information on the structural integrity of the underlying hippocampal networks, analyses on white matter tracts (cingulum bundle) and behavioral responses (rejection rates of syllogisms) were conducted and revealed a negative correlation between cingulum bundle integrity and rejection rate among younger adults (left cingulum bundle: *r*(29) = −0.44, *p* = 0.017; right cingulum bundle: *r*(29) = −0.39, *p* = 0.033), suggesting that younger adults who had higher integrity in the cingulum, rejected the *unbelievable* premises less.

A positive correlation was found between rejection rate of *believable* premises and cingulum integrity among older adults (left cingulum: *r*(29) = 0.38, *p* = 0.038; Table 5), suggesting that older adults who had higher integrity in the left cingulum bundle, rejected the *believable* premises more. No other correlations were found between the integrity of the uncinate fasciculus and performance (all *p*s > 0.05). None of the correlations were significant with RTs.

**Table 5.** Pearson correlation between cingulum bundle and uncinate fasciculus FA values and rejection rate in both age groups

|  | Younger adults |  | Older adults |  |
| --- | --- | --- | --- | --- |
|  | *r* | *df* | *r* | *df* |
| ***Rejection rates (Believable premise)*** |  |  |  |  |
| Right Cingulum | .033 | 29 | .246 | 29 |
| Left Cingulum | -.019 | 29 | **.388** | 29 |
| Right Uncinate Fasciculus | .017 | 29 | .225 | 29 |
| Left Uncinate Fasciculus | .012 | 29 | .352 | 29 |
| ***Rejection rates (Unbelievable premise)*** |  |  |  |  |
| Right Cingulum | **-.396** | 29 | -.075 | 29 |
| Left Cingulum | **-.441** | 29 | .001 | 29 |
| Right Uncinate Fasciculus | .018 | 29 | -.220 | 29 |
| Left Uncinate Fasciculus | .212 | 29 | -.161 | 29 |
| ***Rejection rates (Believable conclusion)*** |  |  |  |  |
| Right Cingulum | -.190 | 29 | .200 | 29 |
| Left Cingulum | -.191 | 29 | .249 | 29 |
| Right Uncinate Fasciculus | .028 | 29 | .207 | 29 |
| Left Uncinate Fasciculus | .088 | 29 | .345 | 29 |
| ***Rejection rates (Unbelievable conclusion)*** |  |  |  |  |
| Right Cingulum | -.208 | 29 | .124 | 29 |
| Left Cingulum | -.266 | 29 | .296 | 29 |
| Right Uncinate Fasciculus | .157 | 29 | -.148 | 29 |
| Left Uncinate Fasciculus | .144 | 29 | -.041 | 29 |

*Note:* Bold numbers indicate significant level less than 0.05.

## Discussion

The present study lends evidence for the age-related differences in logical reasoning and the impact of currently-held beliefs using a syllogism task. First, the whole-brain results from the premise integration stage (second premise) showed that while younger adults recruited a single network for all conditions, older adults’ brain activity was modulated by the believability load of the premise. Our functional connectivity results using the hippocampus as a seed revealed that older adults engaged a hippocampal network for believable premises and this network contributed to higher rejection rates, suggesting more controls over their currently-held beliefs and better logical reasoning performance overall for believable inferences. This network specifically included anterior cingulate and inferior frontal gyrus, regions that are involved in cognitive control and inhibitory control. Furthermore, our structure-function analyses suggested a positive correlation between cingulum bundle structural integrity and rejection rate for believable inferences, that is, the higher the integrity in the cingulum bundle was associated with the higher the rejection rates in believable inferences among older adults. In sum, our results using multimodal imaging; behavioral performance, functional connectivity, and white matter structural integrity together support the compensatory role of hippocampus-prefrontal areas contributing to inhibition of currently-held beliefs during a logical reasoning task among older adults.

### Behavioral findings

Our behavioral results showed a higher rejection rate for unbelievable premises than believable and neutral ones. Higher rates of rejection for unbelievable statements are in line with the existing theories, such as mental model theory, that individuals construct mental models from syllogisms first. If the belief load of the syllogisms conflicts with the mental model, it initiates a search for an alternative model of the premises (Johnson-Laird, 2001; Johnson-Laird, 2010; Johnson-Laird et al., 2015). Our results extend previous studies, which have been mainly focused on the believability load of the conclusion, and suggest that even during the unbelievable premises, the cognitive control might be triggered which leads to differences in logical response.

### Age-related differences during premise integration

To our knowledge, this study is the first to examine neural correlates of belief-content conflict (where the content of premises conflicts with the currently-held belief) in late adulthood and to report the role of IFG and the hippocampus (see the following section) during premise integration stage using a syllogistic reasoning task. Our whole-brain analysis showed that older adults activated several brain areas including left IFG during the premise integration stage, more so for believable and unbelievable conditions than the neutral ones. The importance of the IFG region has been shown in a number of tasks including logical reasoning (Goel & Dolan, 2003; Prado et al., 2011) and cognitive control (Brass et al., 2005; Derrfuss et al., 2005) when tasks are complex and attentional demand is high. There is also neuroimaging evidence that in addition to the inhibitory control (Aron et al., 2014), the IFG is involved in the rehearsal system of working memory (McDermott et al., 2003). During the syllogistic reasoning task, information is needed to be retrieved from memory and currently-held beliefs are required to be inhibited to make sound logical decisions. While previous studies reported the role of IFG during the conclusion stage with various believability load, we have shown that the IFG plays a critical role during the premise integration stage. Our findings take these studies further and suggest that IFG contributes to the inhibition of current beliefs during the premise integration stage in addition to the conclusion stage reported previously. Our results are also in line with findings from using near-infrared spectroscopy method suggesting enhanced IFG activity among older adults when currently-held beliefs are required to be ignored for a sound reasoning decision (Tsujii et al., 2010).

### Age-related differences in the hippocampal functional network during premise integration

A growing body of studies has shown that the hippocampal structural and functional changes contribute to memory and cognitive performance and, thus, are especially important in late adulthood. In addition to the substantial structural changes in the hippocampus and medial temporal cortex volume, which differentiate between healthy and pathological (e.g. AD) aging (Desikan et al., 2006), structural and functional connectivity between the hippocampus and frontal regions go under substantial changes in aging (Fjell & Walhovd, 2010). In our task, the engagement of the hippocampus during premise stage suggests that there is a need to compare the belief content of syllogisms with current beliefs stored in the memory during a logical decision making. Previous studies have suggested that the hippocampus can detect deviant stimuli from their context in the environment (Barbeau et al., 2017; Grunwald et al., 1998), and can detect a mismatch from a novel sequence of events (Garrido et al., 2015). When faced with a logical reasoning task, individuals are required to retrieve semantic knowledge and subsequently, to compare them with assumptions presented at syllogisms. Given the importance of the hippocampus in semantic memory (Manns et al., 2003), our findings offer empirical evidence to the idea that the hippocampus, and its connection with prefrontal areas, are involved in the premise integration stage possibly via retrieval-mediated learning. Our results also suggest that age-related changes in the reasoning might be due to the underlying changes in the hippocampal structure and function. However, further investigation is needed to determine the differential role of the hippocampus and its connection to PFC in various forms of complex reasoning tasks and different stages of reasoning including the conclusion stage.

In our functional connectivity findings, older adults engaged the hippocampus network that included anterior cingulate and inferior frontal gyrus, more for believable premises and this network contributed to rejecting believable assumptions (i.e. higher logical reasoning performance). The engagement of the hippocampus-prefrontal network for believable premises highlights the importance of retrieving semantic associations when the belief load is congruent with currently-held beliefs (Wendelken & Bunge, 2010) and suggests that this network is pivotal in controlling currently-held assumptions and in reaching a logically correct conclusion. The contribution of this network to performance among older adults corroborate the view of engaging a compensatory network by advancing age (for reviews see (Davis et al., 2008; Grady, 2012; Ziaei & Fischer, 2016). Interestingly, a recent study showed that age-related decline in memory-dependent decisions can be diminished by a compensatory network between ventromedial and dorsolateral PFC regions (Lighthall et al., 2014). This over-recruitment of frontal areas during the believable condition in second LV observed in this study, also highlights that functional compensation of PFC regions may be a protective mechanism during the logical decision when rejection of believable assumptions is required.

Another important point concerning the hippocampal activity is that various parts of the hippocampus are involved in rather different tasks. The coordinates for this study were from the posterior part of the hippocampus. Our results are in line with previous reports about the posterior hippocampus to work in concert with regions involved in imagery and perceptual processing to form mental constructions via relational processing (Sheldon & Levine, 2016; Sheldon et al., 2016). Strange and colleagues (1999) also reported the anterior–posterior familiarity gradient, suggesting that an increase in familiarity leads to activation of the posterior hippocampus. Given all categories in the believable condition are familiar categories (e.g., furniture, fruits, animals), different networks might be involved in reasoning about familiar or unfamiliar concepts, i.e. believable vs. unbelievable conditions. Future studies, thus, are needed to distinguish between anterior and posterior divisions of the hippocampus during a logical reasoning task.

### Age-related differences in the structural integrity of cingulum bundle for premise integration

The age-related changes in macrostructural brain properties lead to decreased volumes and thickness in the prefrontal and temporal regions, specifically the prefrontal cortex and hippocampus, which are among well-established findings (Persson et al., 2012). Accumulating evidence has reported the link between the integrity of various white matter pathways and cognitive performances such as response time (Madden et al., 2009); task-switching performance (Gold et al., 2010); working memory (Burianová et al., 2015); motor performance; and problem-solving (Zahr et al., 2009). Specifically, studies have supported the role of the cingulum bundle in tasks associated with visuospatial processing and memory (Davis et al., 2009). Our results are in line with these reports and suggest that, for older adults, FA values derived from the cingulum bundle were positively correlated with the rejection rate of the believable inferences and negatively correlated with the rejection rate of the unbelievable inferences. This finding is in line with our work suggesting the role of the cingulum bundle in the rejection rate for believable statements via engaging the anterior cingulate cortex (Ziaei et al., 2019). This finding suggests that higher integrity in this tract leads to more logically correct decisions that are less bounded by the belief load of the premises, hence, a better logical reasoning performance. Given that this is the first study to investigate the relationship between structural and functional networks in logical reasoning, these results should be considered preliminary and interpreted cautiously. Further investigation is needed to provide conclusive evidence for the role of cingulum in logical reasoning.

### Future directions

It has to be noted that one limitation of this study is the sample size for correlational analyses between cingulum bundle FA values and behavioral measures. Further studies are warranted to replicate the correlational findings in a larger sample size. Additionally, future studies should investigate the role of cortical and subcortical areas in different forms of reasoning such as abductive reasoning. Lastly, it will be informative for future studies to include incorrect responses to the syllogistic reasoning and investigate corresponding brain networks when participants made an incorrect response.

### Conclusion

The primary aim of this study was to investigate the hippocampal-prefrontal functional networks involved in logical reasoning and inhibition of beliefs. For the first time, we have shown that older adults’ brain activity was modulated by the belief load during premise integration stage. Functional connectivity results with the hippocampus revealed that older, but not younger, adults engaged a hippocampal-PFC network which contributed to higher reasoning performance for believable inferences. Additionally, our structure-function connectivity analyses showed that higher cingulum bundle integrity correlated with better logical reasoning performance for believable inferences among older adults’ group. These novel results highlight the importance of the integrity of retrieving semantic information during a logical reasoning task and suggest a compensatory role of anterior prefrontal regions during a reasoning task. This study provides new insights for the relationship between semantic memory, inhibitory control, and logical reasoning in aging, which can be utilized to design appropriate intervention for improving logical reasoning performance.

## Acknowledgment

The authors would like to thank the participants for their time and acknowledge the practical support provided by the imaging staff at the Centre for Advanced Imaging. We also would like to express our appreciation to Ashley York who helped greatly during the data collection stage of this project. This work was supported by funding from the Australian Research Council Science of Learning Special Research Initiative (SR120300015). The authors declare no competing financial interests.

## Notes

### Competing Interest Statement

The authors have declared no competing interest.

