## Supplementary Material for "Role of the hippocampus during logical reasoning and belief bias in aging" (from another version of this preprint: version 1, 2019-11-27)

| **Premise 1** | **Premise 2** | **Conclusion** | **Correct response** | **CoNditioN** | **Proposition type** |
| --- | --- | --- | --- | --- | --- |
| All sodas are drinks | No sodas are soft drink | Therefore: All drinks are soft drink | 2 | U-U | A-E |
| All dralis are soft drink | No dralis are sodas | Therefore: Some soft drink are not sodas | 1 | N-B | A-E |
| All grapefruits are citruses | All citruses are fruits | Therefore: No fruits are grapefruits | 2 | B-U | A-A |
| All fishes are animals | All fishes are moths | Therefore: No animals are moths | 2 | U-U | A-A |
| All ducks are birds | No birds are turtles | Therefore: No ducks are turtles | 1 | B-B | A-E |
| All hammers are mouts | All mouts are snipers | Therefore: All hammers are snipers | 1 | N-U | A-A |
| All lemons are citruses | No citruses are fruits | Therefore: All lemons are fruits | 2 | U-B | A-E |
| All punties are furnitures | All punties are sofas | Therefore: Some furnitures are sofas | 1 | N-B | A-A |
| All sothods are fruits | All sothods are grapefruits | Therefore: All grapefruits are fruits | 2 | N-B | A-A |
| All pistols are weapons | No weapons are furniture | Therefore: All furniture are pistols | 2 | B-U | A-E |
| All punties are fruits | No punties are vegetables | Therefore: All fruits are vegetables | 2 | N-U | A-E |
| All cruises are boats | All cruises are kayaks | Therefore: Some boats are kayaks | 1 | U-B | A-A |
| All sparrows are birds | All birds are animals | Therefore: Some animals are not sparrows | 2 | B-B | A-A |
| All lemons are citruses | All citruses are fruits | Therefore: No fruits are lemons | 2 | B-U | A-A |
| All sodas are drinks | No drinks are soft drink | Therefore: All sodas are soft drink | 2 | B-B | A-E |
| All apples are round fruits | No apples are fruits | Therefore: Some round fruits are not fruits | 1 | U-U | A-E |
| All grapefruits are mesoations | No mesoations are fruits | Therefore: No fruits are grapefruits | 1 | N-U | A-E |
| All lories are goples | All goples are ducks | Therefore: All ducks are lories | 2 | N-U | A-A |
| All blans are animals | No blans are crocodiles | Therefore: All crocodiles are animals | 2 | N-B | A-E |
| All cabbages are vegetables | No vegetables are peaches | Therefore: No cabbages are peaches | 1 | B-B | A-E |
| All round fruits are envenches | All envenches are peaches | Therefore: Some peaches are round fruits | 1 | N-B | A-A |
| All citruses are fruits | No fruits are radishes | Therefore: All citruses are radishes | 2 | B-U | A-E |
| All parrots are animals | No parrots are birds | Therefore: Some animals are not birds | 1 | U-B | A-E |
| All grapefruits are citruses | All grapefruits are lemons | Therefore: No citruses are lemons | 2 | U-U | A-A |
| All apples are fruits | No apples are round fruits | Therefore: All round fruits are fruits | 2 | U-B | A-E |
| All citruses are trinnels | No trinnels are carrots | Therefore: All carrots are citruses | 2 | N-U | A-E |
| All cockatoos are birds | All birds are animals | Therefore: All cockatoos are animals | 1 | B-B | A-A |
| All peaches are round fruits | No peaches are fruits | Therefore: Some round fruits are not fruits | 1 | U-U | A-E |
| All cockatoos are parrots | All cockatoos are moths | Therefore: Some moths are parrots | 1 | U-U | A-A |
| All lories are parrots | All parrots are birds | Therefore: Some birds are not lories | 2 | B-B | A-A |
| All oranges are round fruits | All oranges are peaches | Therefore: All peaches are round fruits | 2 | U-B | A-A |
| All parrots are birds | All parrots are animals | Therefore: No animals are birds | 2 | B-U | A-A |
| All lories are animals | All lories are parrots | Therefore: Some animals are parrots | 1 | B-B | A-A |
| All canaries are birds | All canaries are animals | Therefore: No animals are birds | 2 | B-U | A-A |
| All lemonades are soft drink | All soft drink are beers | Therefore: All lemonades are beers | 1 | U-U | A-A |
| All terrorists are dralys | All dralys are robbers | Therefore: All terrorists are robbers | 1 | N-U | A-A |
| All mammals are bartroods | All bartroods are elephants | Therefore: All elephants are mammals | 2 | N-U | A-A |
| All cockatoos are shigands | No shigands are animals | Therefore: Some animals are not cockatoos | 1 | N-B | A-E |
| All coffees are drinks | All coffees are whiskies | Therefore: Some drinks are whiskies | 1 | U-B | A-A |
| All sodas are drinks | No drinks are soft drink | Therefore: Some sodas are soft drink | 2 | B-B | A-E |
| All pines are trees | No pines are willows | Therefore: All trees are willows | 2 | B-U | A-E |
| All giraffes are curges | All curges are crocodiles | Therefore: No crocodiles are giraffes | 2 | N-B | A-A |
| All sparrows are animals | No sparrows are birds | Therefore: Some animals are not birds | 1 | U-B | A-E |
| All kayaks are boats | No kayaks are cruises | Therefore: All boats are cruises | 2 | B-U | A-E |
| All cupboards are furniture | All cupboards are spears | Therefore: No furniture are spears | 2 | U-B | A-A |
| All parrots are naluals | No naluals are lories | Therefore: All lories are parrots | 2 | N-B | A-E |
| All lemons are brimbers | No brimbers are citruses | Therefore: No citruses are lemons | 1 | N-U | A-E |
| All lemonades are drinks | No lemonades are soft drink | Therefore: All drinks are soft drink | 2 | U-U | A-E |
| All salmons are animals | No salmons are giraffes | Therefore: All giraffes are animals | 2 | B-B | A-E |
| All lemonades are drinks | All drinks are soft drink | Therefore: All lemonades are soft drink | 1 | U-B | A-A |
| All apples are round fruits | No round fruits are fruits | Therefore: No apples are fruits | 1 | U-U | A-E |
| All cockatoos are dralis | No dralis are insects | Therefore: All cockatoos are insects | 2 | N-U | A-E |
| All desks are furniture | All furniture are guns | Therefore: All guns are desks | 2 | U-U | A-A |
| All grapefruits are citruses | No grapefruits are round fruits | Therefore: No citruses are round fruits | 2 | U-B | A-E |
| All lizards are dralis | No dralis are crocodiles | Therefore: No crocodiles are lizards | 1 | N-B | A-E |
| All apples are fruits | No apples are lemons | Therefore: All fruits are lemons | 2 | B-U | A-E |
| All bookcases are furniture | No bookcases are beds | Therefore: Some furniture are not beds | 1 | B-B | A-E |
| All apples are fruits | All apples are round fruits | Therefore: No fruits are round fruits | 2 | B-U | A-A |
| All pistols are mesoations | All mesoations are guns | Therefore: All pistols are guns | 1 | N-B | A-A |
| All pumes are guns | All pumes are desks | Therefore: Some desks are guns | 1 | N-U | A-A |
| All cockatoos are birds | All cockatoos are animals | Therefore: All birds are animals | 2 | B-B | A-A |
| All lemons are citruses | All lemons are fruits | Therefore: No citruses are fruits | 2 | B-U | A-A |
| All oranges are round fruits | No round fruits are fruits | Therefore: All fruits are oranges | 2 | U-U | A-E |
| All moths are trinnels | All trinnels are fishes | Therefore: No fishes are moths | 2 | N-B | A-A |
| All lories are birds | All birds are animals | Therefore: No animals are lories | 2 | B-U | A-A |
| All lories are parrots | No parrots are animals | Therefore: Some animals are not lories | 1 | U-B | A-E |
| All goples are trees | All goples are willows | Therefore: No trees are willows | 2 | N-U | A-A |
| All lories are birds | All birds are snakes | Therefore: All lories are snakes | 1 | U-U | A-A |
| All grapefruits are fruits | All grapefruits are round fruits | Therefore: All round fruits are fruits | 2 | B-B | A-A |
| All desks are furniture | No desks are snipers | Therefore: Some furniture are not snipers | 1 | B-B | A-E |
| All lories are birds | No birds are animals | Therefore: No animals are lories | 1 | U-U | A-E |
| All aschitations are animals | No aschitations are birds | Therefore: All animals are birds | 2 | N-U | A-E |
| All peaches are gruts | No gruts are round fruits | Therefore: All peaches are round fruits | 2 | N-B | A-E |
| All cockatoos are animals | All cockatoos are birds | Therefore: Some animals are birds | 1 | B-B | A-A |
| All sparrows are birds | No birds are animals | Therefore: All sparrows are animals | 2 | U-B | A-E |
| All desks are furniture | All furniture are shovels | Therefore: No desks are shovels | 2 | U-B | A-A |
| All bunges are animals | All bunges are crocodiles | Therefore: Some animals are crocodiles | 1 | N-B | A-A |
| All swords are weapons | All weapons are beds | Therefore: All beds are swords | 2 | U-U | A-A |
| All trops are citruses | No trops are fruits | Therefore: Some citruses are not fruits | 1 | N-U | A-E |
| All apples are fruits | No apples are oranges | Therefore: All fruits are oranges | 2 | B-U | A-E |
| All oranges are citruses | No citruses are fruits | Therefore: All fruits are oranges | 2 | U-U | A-E |
| All sparrows are birds | All birds are lizards | Therefore: No lizards are sparrows | 2 | U-B | A-A |
| All renes are weapons | No renes are hammers | Therefore: Some weapons are not hammers | 1 | N-B | A-E |
| All papes are fruits | All papes are lemons | Therefore: No fruits are lemons | 2 | N-U | A-A |
| All goples are round fruits | No goples are fruits | Therefore: Some round fruits are not fruits | 1 | N-U | A-E |
| All apples are round fruits | All round fruits are fruits | Therefore: All apples are fruits | 1 | B-B | A-A |
| All cokes are drinks | All drinks are soft drink | Therefore: All cokes are soft drink | 1 | U-B | A-A |
| All desks are furnitures | No furnitures are spears | Therefore: All desks are spears | 2 | B-U | A-E |
| All sparrows are birds | All birds are animals | Therefore: No animals are sparrows | 2 | B-U | A-A |
| All cockatoos are parrots | All cockatoos are crocodiles | Therefore: Some crocodiles are parrots | 1 | U-U | A-A |
| All oranges are citruses | No citruses are fruits | Therefore: Some fruits are not oranges | 1 | U-B | A-E |
| All spaves are furniture | No spaves are snipers | Therefore: No furniture are snipers | 2 | N-B | A-E |
| All striffs are soft drink | All striffs are teas | Therefore: Some soft drink are teas | 1 | N-U | A-A |
| All lemons are round fruits | No round fruits are vegetables | Therefore: All lemons are vegetables | 2 | B-U | A-E |
| All grapefruits are fruits | No grapefruits are parsnips | Therefore: No fruits are parsnips | 2 | B-B | A-E |
| All duyumences are guns | All duyumences are sofas | Therefore: No guns are sofas | 2 | N-B | A-A |

Responses were either follow (1) or not follow (2). Conditions coded based on the belief load of the premise and conclusion. First letter in the condition column represents the believability of the premise (B = believable, N = neutral, U = unbelievable) and second letter represents the belief load of the conclusion (B = believable, U = unbelievable). Two types for the proposition were used: A refers to all and E refers to No. First letter corresponds to the second premise’s and second letter corresponds to the conclusion’s proposition type.

**Syllogisms structure**

Logical arguments in this study were in the form of standard syllogisms (Table 1) and included three statements: two premises and one conclusion. The subject and the predicate of a premise was formed by arbitrary sets (e.g., dogs, mammals, furniture). The two premises had exactly one set in common that may appear in either the subject or the predicate in either of the premises (${Set}_{2}$ in Table 1). Hence, the two premises involved exactly three sets (${Set}_{1}$, ${Set}_{2}$, and ${Set}_{3}$ in the example in Table 1), two of which are uniquely used in each premise and one which is used in both. The conclusion of a syllogism provides a statement about the sets that appear uniquely in premises (${Set}_{1}$ and ${Set}_{3}$ in the example). A conclusion “follows” from the premises if the premises provide conclusive evidence to support it. Otherwise, the conclusion “does not follow” from the premises, either because the conclusion is wrong given the premises, or is not completely supported by the premises.

*Table 1. Form of syllogisms, quantifiers, subjects, copula, and predicate used in the experimental design*

|  | Quantifier | Subject | Copula | Predicate |
| --- | --- | --- | --- | --- |
| Premise 1 | All | $\mathrm{Set}_{1}$ | is | $\mathrm{Set}_{2}$ |
| Premise 2 | No | $\mathrm{Set}_{2}$ | is | $\mathrm{Set}_{3}$ |
| Conclusion | Some | $\mathrm{Set}_{3}$ | is not | $\mathrm{Set}_{1}$ |

Additionally, in a syllogism, each statement includes a quantifier (All, No, and Some) and a copula (Is or Is not). We used propositions A (All, is) and E (No, is) in all premises (counter balanced across all runs). Given that proposition E simplifies the reasoning task substantially, we avoided the use of proposition E in both premises. For the conclusions, however, we used the appropriate propositions required to ensure that conclusions that followed or did not follow the premises were balanced.

The order of the sets and predicate of the premises were balanced. In this experiment, we used two types of ordering: i) the common set belonged to the subject of premise 1 and the subject of premise 2, and ii) the common set belonged to the predicate of premise 1 and the subject of premise 2. The conclusion takes the unique sets from each premise and combines them through a proposition. The logical reasoning task was to decide if the generated statement follows from the premises or not. Whether the subject of the conclusion was drawn from the first or second premise was also counterbalanced.
